# The division defect of a *Bacillus subtilis minD noc* double mutant can be suppressed by Spx-dependent and Spx-independent mechanisms

**DOI:** 10.1101/2021.05.05.442877

**Authors:** Yuanchen Yu, Felix Dempwolff, Reid T. Oshiro, Frederico J. Gueiros-Filho, Stephen C. Jacobson, Daniel B. Kearns

## Abstract

During growth, bacteria increase in size and divide. Division is initiated by the formation of the Z-ring, an intense ring-like cytoskeletal structure formed by treadmilling protofilaments of the tubulin homolog FtsZ. FtsZ localization is thought to be controlled by the Min and Noc systems, and here, we explore why cell division fails at high temperature when the Min and Noc systems are simultaneously mutated. Microfluidic analysis of a *minD noc* double mutant indicated that FtsZ formed proto-Z-rings at periodic inter-chromosome locations but that the rings failed to mature and become functional. Extragenic suppressor analysis indicated that a variety of mutations restored high temperature growth to the *minD noc* double mutant, and while many were likely pleiotropic, others implicated the proteolysis of the transcription factor Spx. Further analysis indicated that a Spx-dependent pathway activated the expression of ZapA, a protein that primarily compensates for the absence of Noc. Additionally, an Spx-independent pathway increased the activity of the divisome to reduce the length of the cytokinetic period. Finally, we provide evidence of an as-yet-unidentified protein that is activated by Spx and governs the frequency of polar division and minicell formation.

**IMPORTANCE:** Bacteria must properly position the location of the cell division machinery in order to grow, divide, and ensure each daughter cell receives one copy of the chromosome. In *B. subtilis*, cell division site selection is thought to depend on two systems called Min and Noc, and while neither is individually essential, cells fail to grow at high temperature when both are mutated. Here, we show that cell division fails in the absence of Min and Noc, not due to a defect in FtsZ localization, but rather a failure in the maturation of the cell division machinery. To understand what happens when the division machinery fails to mature, suppressor mutations that bypass the need for Min, Noc, or both were selected. Some of the mutants activated the Spx stress response pathway while others appeared to directly enhance divisome activity.

## INTRODUCTION

Most bacteria are surrounded by a semi-elastic cell wall made of peptidoglycan that protects the cell membrane from hyperexpansion due to the osmotic pressure of the cytoplasm. When cells divide, peptidoglycan growth must be redirected from synthesis that expands the cell perimeter to synthesis that bisects the mother cell into two daughters.^1^ Ultimately, cell division requires two critical events. First, the cell must identify a site for division that falls between replicated chromosomes such that upon completion, each daughter would receive one full copy of the genome. ^2, 3^ Second, a large complex of proteins known as the divisome must be recruited to the site of cell division and, once activated, synthesizes peptidoglycan to constrict the cell envelope and ultimately complete cytokinesis.^4, 5^ Both events are mediated by FtsZ, a protein that marks the site of cell division and serves as a scaffold for the recruitment of the divisome complex.^6^

FtsZ is a GTP binding and hydrolyzing homolog of eukaryotic tubulin that forms protofilament polymers inside the cell.^7–11^ Early work indicated that FtsZ localized as a ring and was the earliest component of the divisome that recruited subsequent proteins by direct interaction.^12–15^ Later work indicated that localization of FtsZ was dynamic as rings formed at midcell, increased in intensity, constricted in diameter and ultimately, disappeared after completion of cytokinesis.^16–20^ FtsZ dynamism includes, but is not limited to, the phenomenon of treadmilling where each individual monomer is stationary within a protofilament, but the profilament as a whole appears to travel as new monomers are added to one end and lost from the other.^21–24^ FtsZ dynamism is critical for function not only to reposition the FtsZ ring for each round of division but may also play a role in cytokinesis as protofilaments are in constant motion circumnavigating the division plane.^25–27^ Dynamic control of FtsZ localization is thought to be essential in the process of cell division site selection, symmetrical division, and cell shape control.^28, 29^

One system that controls the dynamic localization of FtsZ is the Min system.^30^ Not all bacteria encode Min, but those that do tend to encode the two most central components, MinC and MinD.^31^ MinD is a membrane-associated protein that binds to and activates MinC.^32–35^ MinC in turn binds to and antagonizes FtsZ.^36–39^ Localization of MinC and MinD is topologically restricted by various mechanisms depending on the organism. In *B. subtilis*, MinC and MinD are restricted to the cell poles by the multipass transmembrane protein MinJ and the scaffolding network of DivIVA. ^40–45^ Polar localization is related to Min function, and in the case of *B. subtilis*, the FtsZ ring remains after division is completed and requires the Min system for depolymerization such that in the absence of Min, FtsZ rings persist at the polar location indefinitely.^46–51^ Constitutive maintenance of multiple Z-rings and the inability to recycle FtsZ monomers puts an increased emphasis on de novo FtsZ synthesis for division, and competition between the multiple rings for FtsZ recruitment results in elongated cells.^51–54^ Finally, recruitment of FtsZ to the persistent polar FtsZ rings eventually gives rise to polar division events and the generation of anucleoid minicells.^47, 51, 55^ While cells mutated for the Min system do not suffer a growth rate defect, Min is nonetheless important for FtsZ redistribution and the cell size control.

Another factor that controls the dynamic localization of FtsZ is the phenomenon of nucleoid occlusion that prevents FtsZ localization in the vicinity of the nucleoid.^56–61^ In *B. subtilis*, nucleoid occlusion is mediated, at least in part, by the DNA-binding Noc protein. ^62, 63^ Noc binds to many sites across the chromosome and also encodes an amphipathic helix that targets/anchors Noc to the membrane.^63, 64^ Consistent with a role in nucleoid occlusion, decondensed spiral-like FtsZ intermediates are observed in close proximity to the chromosome when Noc is mutated.^62^ Recent microfluidic analysis indicated that the decondensed FtsZ filaments originated as a subpopulation of FtsZ that departed the cytokinetic ring and migrated along the membrane to the future site of cell division.^65^ Thus, multiple points of Noc recruitment of DNA to the membrane is thought to act as a passive palisade that occludes FtsZ migration over the chromosome and thereby corrals FtsZ between nucleoids. Cells mutated for the Noc system do not suffer a growth rate defect but Noc is nonetheless important for restricting FtsZ migration and concentrating FtsZ to improve division efficiency.

While neither the Min system nor the Noc system is essential when disrupted on their own, double mutants defective in both systems simultaneously are synthetically sick.^62, 66^ Here, we show that a *minD noc* double mutant has a temperature-sensitive synthetic lethal phenotype, and we explore the conditional essentiality by monitoring double mutants in microfluidic growth channels and by isolating mutants that restore high temperature growth. We find that when Noc is depleted in a *minD* mutant, treadmilling FtsZ protofilaments leave the cytokinetic ring and localize to inter-nucleoid spaces, but the FtsZ rings are defective in maturation as they fail to reach sufficient concentration to induce cytokinesis. Suppressor mutations that restored high temperature growth to the *minD noc* double mutant were found in a variety of genes, and we focused on those genes involved in the proteolytic regulation of the global stress response transcription factor Spx. We found that elevated Spx levels increased the expression of the FtsZ- crosslinking protein ZapA, a condition that primarily masks the absence of Noc. We also found that ClpX inhibited another pathway that is Spx-independent and acts downstream of FtsZ localization, likely at the level of the divisome, to increase the rate of cytokinesis.

## RESULTS

### The FtsZ ring localizes normally but fails to mature in the absence of MinD and Noc

The Noc protein was serendipitously discovered when mutations in the *noc* gene were found to be synthetically sick with mutations in the *min* system incubated under growth conditions of 30°C.^62, 66^ Previous work on the double mutant was conducted in domesticated laboratory strains, and to confirm that the growth defect was not dependent on genetic background, we attempted to reproduce the phenotype in the ancestral *B. subtilis* strain. A *minD noc* double mutant was successfully generated and grew like the wild type on plates and in liquid media incubated at 30°C. Due to the absence of a growth defect under reported conditions, we wondered whether the phenotype was temperature sensitive. To assess temperature sensitivity, the wild type, both single mutants, and the double mutant were serially diluted, spotted on LB agar plates separately, and incubated at four different temperatures: 22°C, 30°C, 37°C, and 42°C (**Fig 1A, Fig S1A**). Whereas wild type, *minD*, and *noc* single mutants exhibited similar viability at all temperatures tested, the number of colony forming units in the *minD noc* double mutant was dramatically reduced at 37°C and 42°C **(Fig 1A, Fig S1A**). We conclude that the ancestral strain is more tolerant than laboratory strains for the *minD noc* double mutant, and we further conclude that the combination of the two mutations is synthetic lethal at high temperatures.

**Figure 1:**
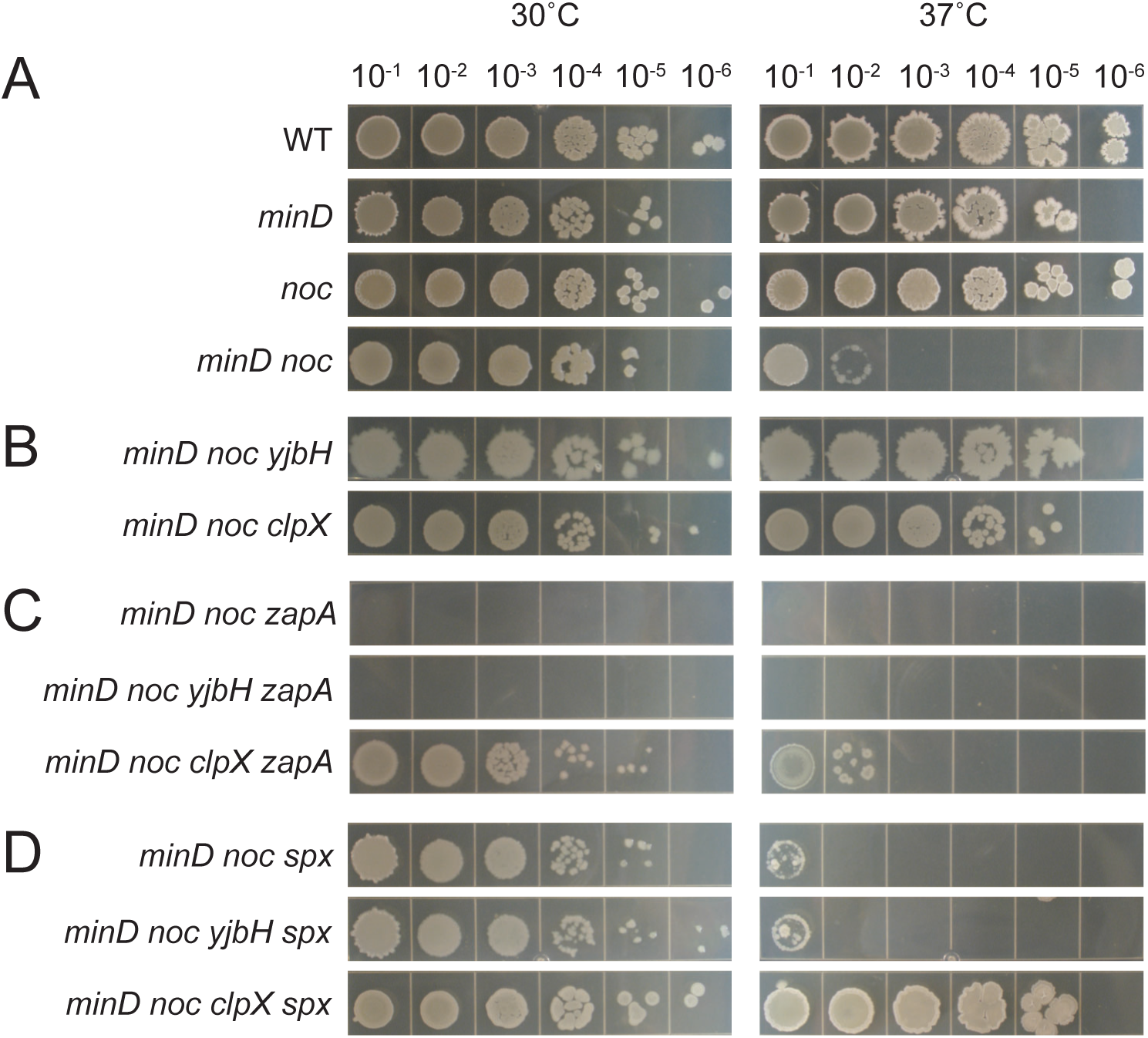
*yjbH* restores high-temperature viability to the *minD noc* double mutant through *spx*. Dilution plating of A) the wild type (DK1042), *minD* (DK7744), *noc* (DK443), *minD noc* (DK7298), B) *minD noc yjbH* (DK8811), *minD noc clpX* (DK8369), C) *minD noc zapA* (DK8800), *minD noc yjbH zapA* (DK8851), *minD noc clpX zapA* (DK8824), and D) *minD noc spx* (DK8767), *minD noc yjbH spx* (DK8849), *minD noc clpX spx* (DK8738) at 30°C and 37°C. Exponential growing liquid culture were diluted to OD_600_ ≈1 and then serially diluted to 10^-1^, 10^-2^, 10^-3^, 10^-4^, 10^- 5^, and 10^-6^. 10 µL of cells were spotted for each concentration. Additional temperatures of 22°C and 42°C are included in supplemental **Fig S1**.

To explore why cells failed to grow at high temperature in the absence of both the Min and Noc systems, we generated a conditional double mutant in which *noc* gene was expressed from an IPTG-inducible promoter and inserted at an ectopic site in the chromosome (*minD noc amyE::P_spank_-noc*). To determine functionality, the IPTG-inducible *noc* construct was introduced at low temperature to a *minD noc* double mutant background containing both the cytoplasmic mRFPmars and mNeongreen-FtsZ fusion reporters. The IPTG-inducible construct appeared to be functional and capable of complementing a *noc* mutation as the strain resembled the *minD* mutant when grown in broth at 37°C with ≥ 0.1 mM IPTG, but showed increased filamentation at lower levels of IPTG induction (**Fig S2A**). Next, the strain was introduced to the microfluidic channels and grown at 37°C with 0.1 mM IPTG induction for three hours after which the IPTG was washed out and replaced with IPTG-deficient media. *B. subtilis* divides by plate formation (septation) with subsequent cell separation, and thus, cell division events were conservatively defined as a local 20% decrease in mRFPmars cell body intensity.^51, 65^ Prior to IPTG washout, cells of the *minD noc* double mutant induced for *noc* grew in a manner similar to that reported for a *minD* mutant alone such that cells were elongated, and frequent minicells were observed (**Fig 2A**).^51^ Over the next 5-6 hours, the cells grew but gradually stopped dividing until the entire population became filamentous and ultimately lysed **(Fig 2B**) (**Movie S1**). We conclude that *noc* depletion in a *minD* mutant background resulted in a cell division defect and cell death consistent with synthetic lethality.

**Figure 2:**
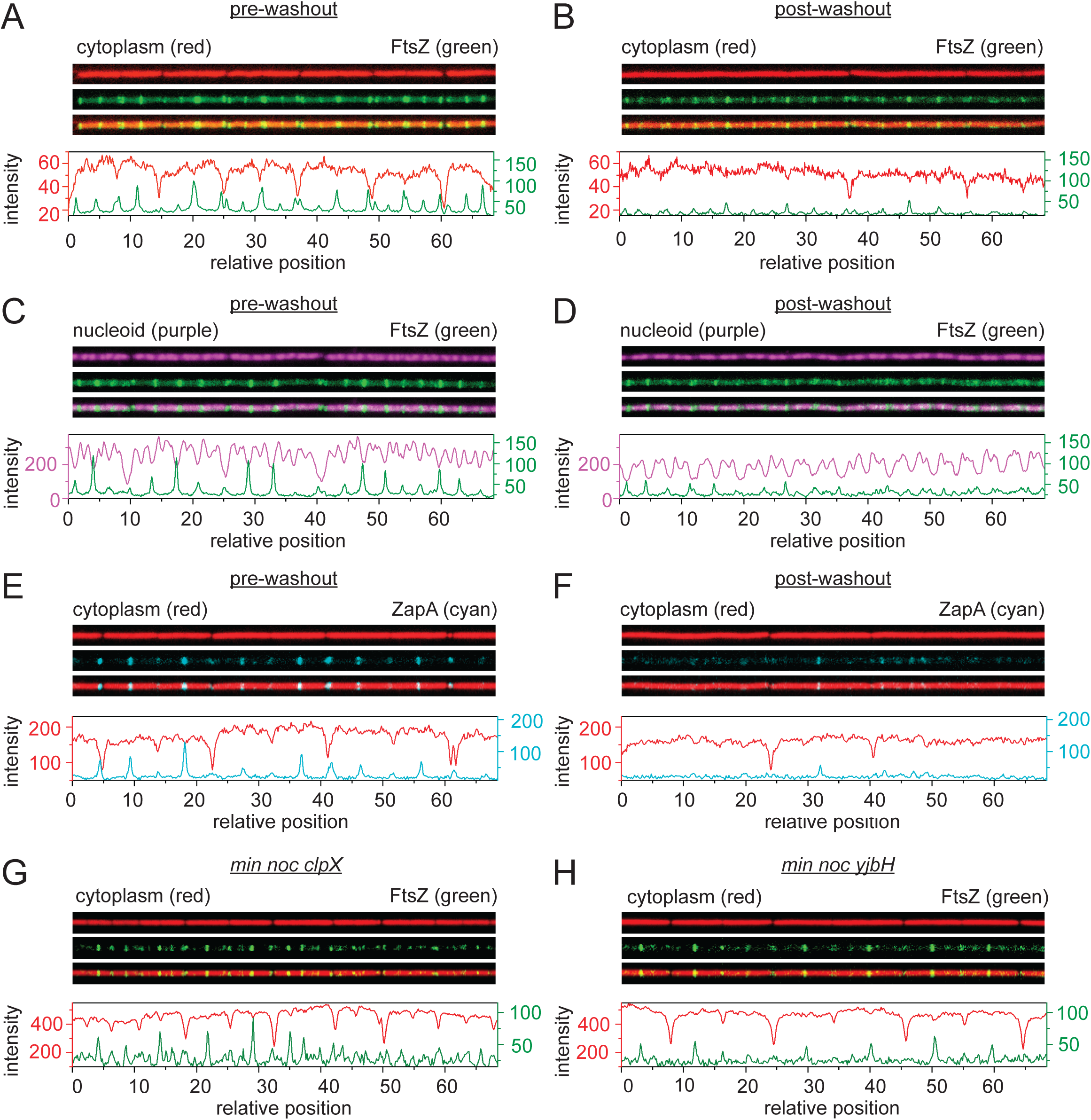
Microfluidic analysis of growth, division, and chromosome segregation in *min noc* double mutants. Time-lapsed fluorescence imaging of a microfluidic channel was performed for the *min noc* (*P_spank_-noc*) mutants (A-F), *minD noc clpX* mutant (G), and *minD noc yjbH* mutant (H) growing at steady state. A and B) Fluorescence microscopy of a *min noc* (*P_spank_-noc*) mutant (DK6375) growing with 0.1 mM IPTG (A) and after washout with media lacking IPTG (B). Cytoplasmic mRFPmars false colored red (top), mNeongreen-FtsZ false colored green (middle), and an overlay of the two colors (bottom). Graphs are a quantitative analysis of mRFPmars fluorescence intensity (red line) and mNeongreen fluorescence intensity (green line) to match the fluorescence images immediately above. C and D) Fluorescence images of a *min noc* (*P_spank_- noc*) mutant (DK6962) growing with 0.1 mM IPTG (C) and after washout with media lacking IPTG (D). Chromosomes (HBsu-mCherry) false colored purple (top), mNeongreen-FtsZ false colored green (middle), and an overlay of the two colors (bottom). Graphs are a quantitative analysis of mCherry fluorescence intensity (purple) and mNeongreen fluorescence intensity (green) to match the fluorescence images immediately above. E and F) Fluorescence images of a *min noc* (*P_spank_- noc*) mutant (DK8394) growing with 0.1 mM IPTG (E) and after washout with media lacking IPTG (F). Cytoplasmic mRFPmars false colored red (top), ZapA-mNeongreen false colored cyan (middle), and an overlay of the two colors (bottom). Graphs are a quantitative analysis of mRFPmars fluorescence intensity (red) and mNeongreen fluorescence intensity (cyan) to match the fluorescence images immediately above. G) Fluorescence images of a *min noc clpX* mutant (DK8390) expressing cytoplasmic mCherry protein (falsely colored red; top) and mNeongreen- FtsZ (falsely colored green; middle) with an overlay of the two (bottom). Graphs are a quantitative analysis of mCherry fluorescence intensity (red) and mNeongreen fluorescence intensity (green) to match the fluorescence images immediately above. H) Fluorescence images of a *min noc yjbH* mutant (DK8817) expressing cytoplasmic mCherry protein (falsely colored red; top) and mNeongreen-FtsZ (falsely colored green; middle) with an overlay of the two (bottom). Graphs are a quantitative analysis of mCherry fluorescence intensity (red) and mNeongreen fluorescence intensity (green) to match the fluorescence images immediately above. For all the panels, two different Y-axes are used. The left axis corresponds to the mRFP mars (red), HBsu-mCherry (purple), or mCherry signals (red) and the right axis corresponds to the mNeongreen-FtsZ (green) or ZapA-mNeongreen signal (cyan). All images are reproduced at the same magnification.

Cells might fail to divide when MinD is mutated and Noc is depleted if FtsZ failed to localize at inter-chromosome locations. Prior to IPTG washout, periodic foci of mNeongreen-FtsZ fluorescence were observed either at the midcell, to mark the future site of cell division, or at the cell pole as a remnant of cytokinesis (**Fig 2A**).^51^ After washout, FtsZ still localized as periodic foci but with a reduced fluorescence intensity (**Fig 2B**) (**Movie S2**). Moreover, FtsZ peak fluorescence still formed in between regularly-spaced chromosomes labeled by the expression of mCherry fused to the nucleoid binding protein HBsu both prior to (**Fig 2C**) and after IPTG washout (**Fig 2D**).^67, 68^ Thus, the *minD noc* double mutant fails to divide, not because the FtsZ fails to be periodically enriched in inter-chromosome spaces, but because the Z-rings that nucleate successfully fail to mature to wild type intensity.

One reason that FtsZ might not mature into an intense ring is if the total amount of FtsZ was diminished in the absence of Min and Noc such that the pool of monomers was insufficient. Quantitative analysis of the fluorescence microscopy data, however, indicated that the fluorescence intensity per cell of the *minD noc* double mutant prior to IPTG washout was the same as the fluorescence intensity over the same length interval after washout (**Fig 3A, left**). Thus, FtsZ levels appeared to be constant. In a parallel approach, Western blot analysis with an antibody raised against FtsZ and a separate antibody against the constitutive control protein SigA was performed on the wild type and on the *minD noc (P_spank_-noc)* double mutant in which the *noc* construct was induced with various amounts of IPTG. The amount of FtsZ remained constant relative to the amount of SigA in both strains regardless of the level of IPTG induction **(Fig 3B**). Based on the results of both the cell biological and biochemical analyses, we conclude that the inability to form Z-rings in the absence of MinD and Noc was not due to a reduction in FtsZ levels.

**Figure 3:**
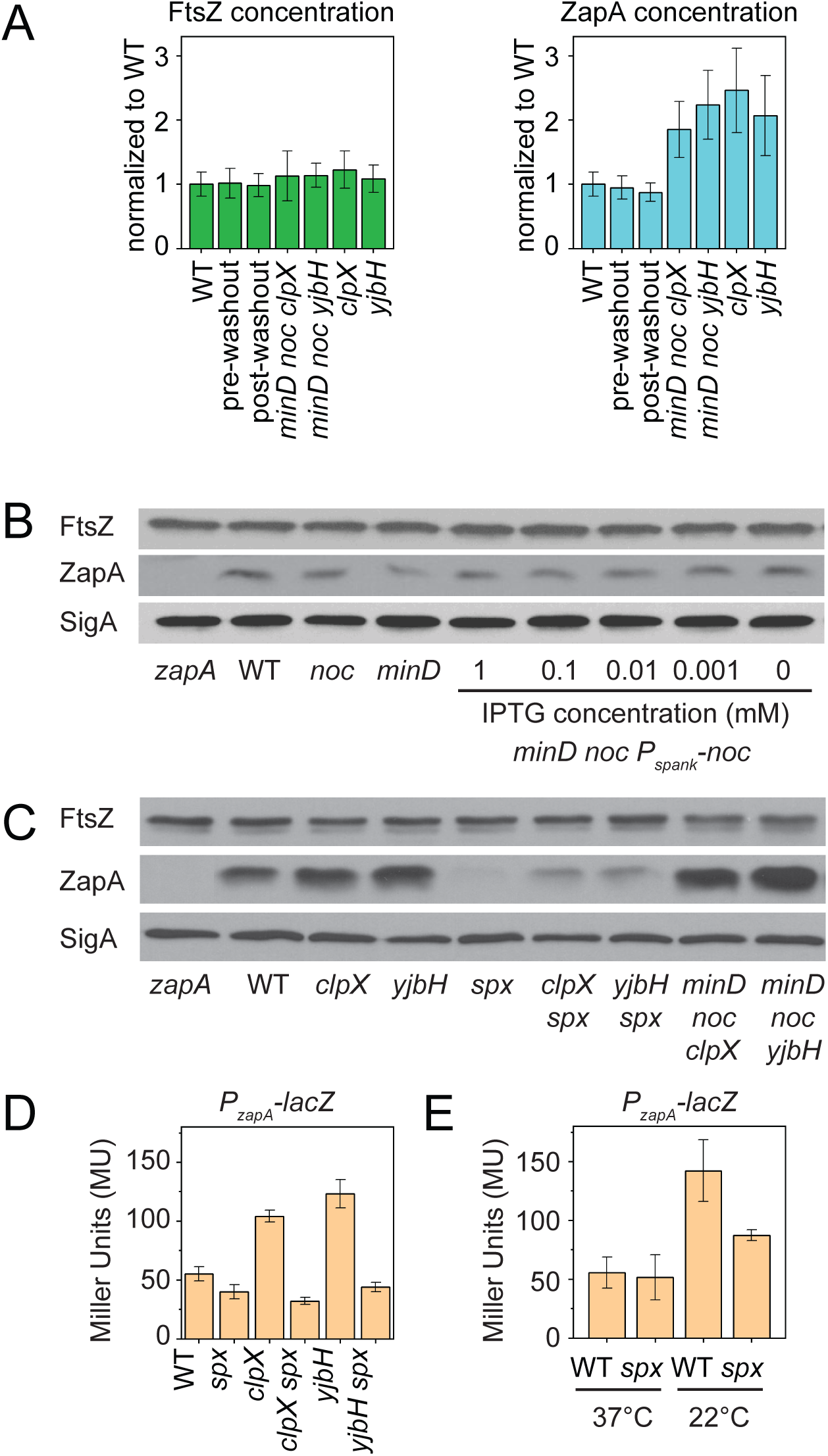
ClpX and YjbH regulate *zapA* expression through Spx. A) Left panel: bar graph of the average mNeongreen-FtsZ fluorescence intensity in individual cells of the wild type (WT, DK8162), *minD noc* (*P_spank_-noc*) (DK6375) mutant growth with 0.1 mM IPTG (pre-washout) and without IPTG (post washout), *minD noc clpX* (DK8390), *minD noc yjbH* (DK8817), *clpX* (DK7320) and *yjbH* (DK8741). Right panel: bar graph of the average ZapA-mNeongreen fluorescence of the wild type (WT, DK8138), *minD noc* (*P_spank_-noc*) (DK8394) mutant growth with 0.1 mM IPTG (pre-washout) and without IPTG (post washout), *minD noc clpX* (DK8387), *minD noc yjbH* (DK8749), *clpX* (DK8147) and *yjbH* (DK8742). Fluorescence intensity of each strain was normalized with the wild type. Error bars are the standard deviations of three replicates. B) Western blot analysis of *B. subtilis* cell lysates of *zapA* (DK8174), wild type (DK1042), *noc* (DK443), *minD* (DK7744), *minD noc* (*P_spank_-noc*) (DK7369) grown in the presence of the amount of IPTG, separately probed with anti-FtsZ primary antibody (top), anti-ZapA primary antibody (middle), and anti-SigA primary antibody (bottom). C) Western blot analysis of *B. subtilis* cell lysates of *zapA* (DK8174), wild type (DK1042), *clpX* (DK5565), *yjbH* (DS8875), *spx* (DK6677), *clpX spx* (DK8880), *yjbH spx* (DK8881), *min noc clpX* (DK8369), and *minD noc yjbH* (DK8811), separately probed with anti-FtsZ primary antibody (top), anti-ZapA primary antibody (middle), and anti-SigA primary antibody (bottom). D) β-Galactosidase assays of *P_zapA_-lacZ* transcriptional fusions for the wild type (DK8591), *spx* (DK8627), *clpX* (DK8628), *clpX spx* (DK8631), *yjbH* (DK8883), *yjbH spx* (DK8890) at 37°C. Error bars are the standard deviations of three replicates. E) β-Galactosidase assays of *P_zapA_-lacZ* transcriptional fusions for the wild type (DK8591) and *spx* (DK8627) grown at either 37°C or at room temperature (22°C) as indicated. Error bars are the standard deviations of three replicates.

Another reason that cells might fail to form intense FtsZ-rings is a failure in Z-ring condensation. Condensation of FtsZ rings appears to be mediated, at least in part, by the FtsZ-associated protein ZapA that binds to and crosslinks FtsZ protofilaments.^20, 69, 70^ To monitor ZapA dynamics, mNeongreen was fused to ZapA and monitored in a *minD noc* (*P_spank_-noc*) mutant background both before and after IPTG washout at 37°C. Prior to washout, ZapA-mNeongreen fluorescence was observed both at the future midcell and at the poles in a pattern reminiscent of that observed for FtsZ (**Fig 2E**). After washout, the ZapA-mNeongreen became more diffuse, like that observed with mNeongreen-FtsZ (**Fig 2F**). Also similar to FtsZ, quantitative analysis indicated that ZapA-mNeongreen levels remained constant after Noc depletion in microfluidic channels (**Fig 3B**), and Western blot analysis with antibodies raised against ZapA protein (**Fig 3C**) indicated that ZapA levels were constant regardless of the level of Noc induction. Unlike mNeongreen-FtsZ, however, ZapA-mNeongreen did not display periodic foci after Noc depletion, but rather only formed intense rings at locations where cell division would eventually occur (**Fig 2F**, **Movie S3**). We conclude that ZapA and FtsZ are maintained at wild type levels in the absence of MinD and Noc, and that ZapA localization is an indicator of mature Z-rings that promote cytokinesis. We further conclude that cells die in the absence of Min and Noc because Z-rings fail to mature.

### An increase in FtsZ ring condensation overrides the absence of Noc in the *minD noc* double mutant

To gain insight into FtsZ-ring maturation, a forward genetic screen was performed to identify genes, which when mutated could permit growth of the *minD noc* double mutant at high temperature. A transposon mutagenesis system was introduced and propagated in the *minD noc* double mutant background, and the resulting transposants from seven parallel mutagenesis experiments were plated at the non-permissive 42°C temperature.^71^ Mutant colonies were isolated from the pool grown at high temperature with a net frequency of approximately 1 x 10^-3^, or 1 in 1000 transposants. Two candidates were clonally isolated from each mutagenesis pool and backcrossed to a *minD noc* double mutant parental background at the permissive temperature. In each case the resulting backcrossed transposon conferred the ability to restore growth at 42°C upon subsequent culturing confirming genetic linkage between the insertion and the phenotype. The location of each of the fourteen transposon insertions was identified by inverse PCR and sequencing (**Table 1**).

**Table 1:**
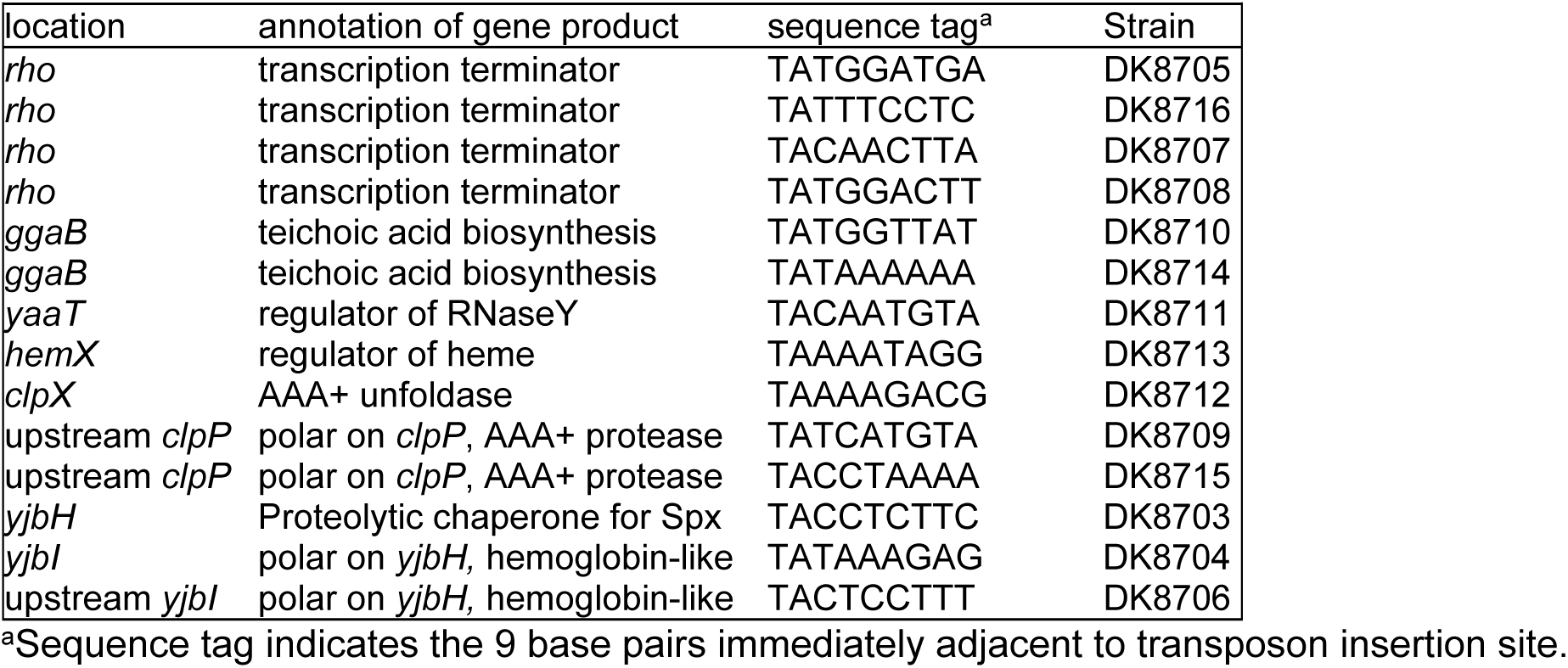
Transposon insertions that restored high temperature growth to a minD noc double mutant.

Eight of the transposons disrupted a variety of genes that could give rise to pleiotropic effects. For example, four of the transposons disrupted the *rho* gene encoding the transcriptional terminator protein Rho, suggesting that RNA anti-termination may be important for growth in the absence of MinD and Noc.^72, 73^ One transposon insertion disrupted the *yaaT* gene, encoding YaaT (RicT), involved in the regulation of Spo0A phosphorylation and RNase Y.^74–77^ Two other insertions were found in the *ggaB* gene, encoding GgaB, a protein involved in teichoic acid biosynthesis,^78^ and one other insertion disrupted *hemX*, encoding HemX, a regulator of HemA- dependent reduction of glutamyl-tRNA.^79^ The six remaining strains, however, were disrupted for genes related to regulatory proteolysis. In particular, three disrupted either the *clpX* gene, encoding an ATP-dependent unfoldase for the ClpP protease,^80, 81^ or inserted upstream of the gene encoding ClpP, such that the amount of ClpP might be reduced by polarity downstream of the insertion site. Importantly, three other transposons either directly disrupted, or were upstream of and likely polar on, the expression of the proteolytic adaptor protein YjbH, that directs ClpXP-mediated degradation of the global stress response transcription factor Spx. ^82–86^ While one or more mechanisms of suppression may be at work, we focused our attention on the mutants disrupted for the ClpXP/YjbH system as their effects could likely be attributed to the Spx protein.

Mutation of either ClpX or YjbH rescued high temperature growth (**Fig 1B**) and efficient cell division to the *minD noc* double mutant background by restoring the formation of intense rings of FtsZ (**Fig 2G,H**). One way in which growth might be improved in the absence of MinD and Noc is if the suppressor mutations in *clpX* and/or *yjbH* increased expression of FtsZ.^62^ The fluorescent intensity of mNeongreen-FtsZ did not appear to increase, however (**Fig 3A, left**), and Western blot analysis indicated that FtsZ protein levels were similar to wild type in either suppressor background (**Fig 3C**). We conclude that growth rescue was not correlated with an increase in FtsZ. Another way in which growth might be improved is if the suppressors increased expression of ZapA, a protein which when overexpressed has been shown to compensate for the absence of Noc.^65^ Fluorescence intensity of ZapA-mNeongreen was two-fold higher (**Fig 3A, right**), and Western blot analysis indicated that transposon insertions in *clpX* and *yjbH* both increased ZapA protein levels relative to wild type (**Fig 3C**). Moreover, mutation of *zapA* exacerbated the temperature sensitive growth phenotype of the *minD noc* double mutant and abolished the ability to rescue growth at 37°C when either *yjbH* or *clpX* were also mutated (**Fig 1C**). We conclude that high-temperature growth rescue in the *minD noc* double mutant by mutation of either *clpX* or *yjbH* was correlated with an increase in ZapA expression and that ZapA was necessary for suppression.

To determine whether an increase in ZapA levels was sufficient to restore high temperature growth in the absence of MinD and Noc, an IPTG-inducible copy of *zapA* was introduced to *a minD noc* double mutant background that also encoded mNeongreen-FtsZ and constitutive cytoplasmic mRFPmars. The resulting strain grew at both 30°C and 37°C even in the absence of IPTG indicating that the low basal level expression from the IPTG-inducible promoter was sufficient to promote growth (**Fig 4A**). At levels up to 0.001 mM IPTG, fluorescence microscopy revealed cells that appeared elongated with diffuse FtsZ fluorescence resembling the strain when no IPTG was added (**Fig 4A**). At 0.01 mM, IPTG however, cell length decreased, FtsZ formed multiple rings per cell, and minicells were observed, perhaps suggesting that the *noc* mutation was being masked thereby revealing a *minD*-like phenotype (**Fig 4A**). As IPTG levels increased to 0.03 mM, excess ZapA became inhibitory, and cells appeared elongated once more (**Fig 4A**). At still higher levels of IPTG, growth was inhibited entirely. Thus, 0.01 mM IPTG was the optimum concentration to bypass the *minD noc* double mutant, and quantitative microfluidic analysis supported that the cell dimensions, FtsZ localization pattern, and minicell frequency resembled that of cells mutated for *minD* alone (**Fig 4B**). We conclude that titrated overexpression of ZapA was sufficient to rescue growth to the *minD noc* double mutant and likely did so by compensating for the absence of Noc.

**Figure 4:**
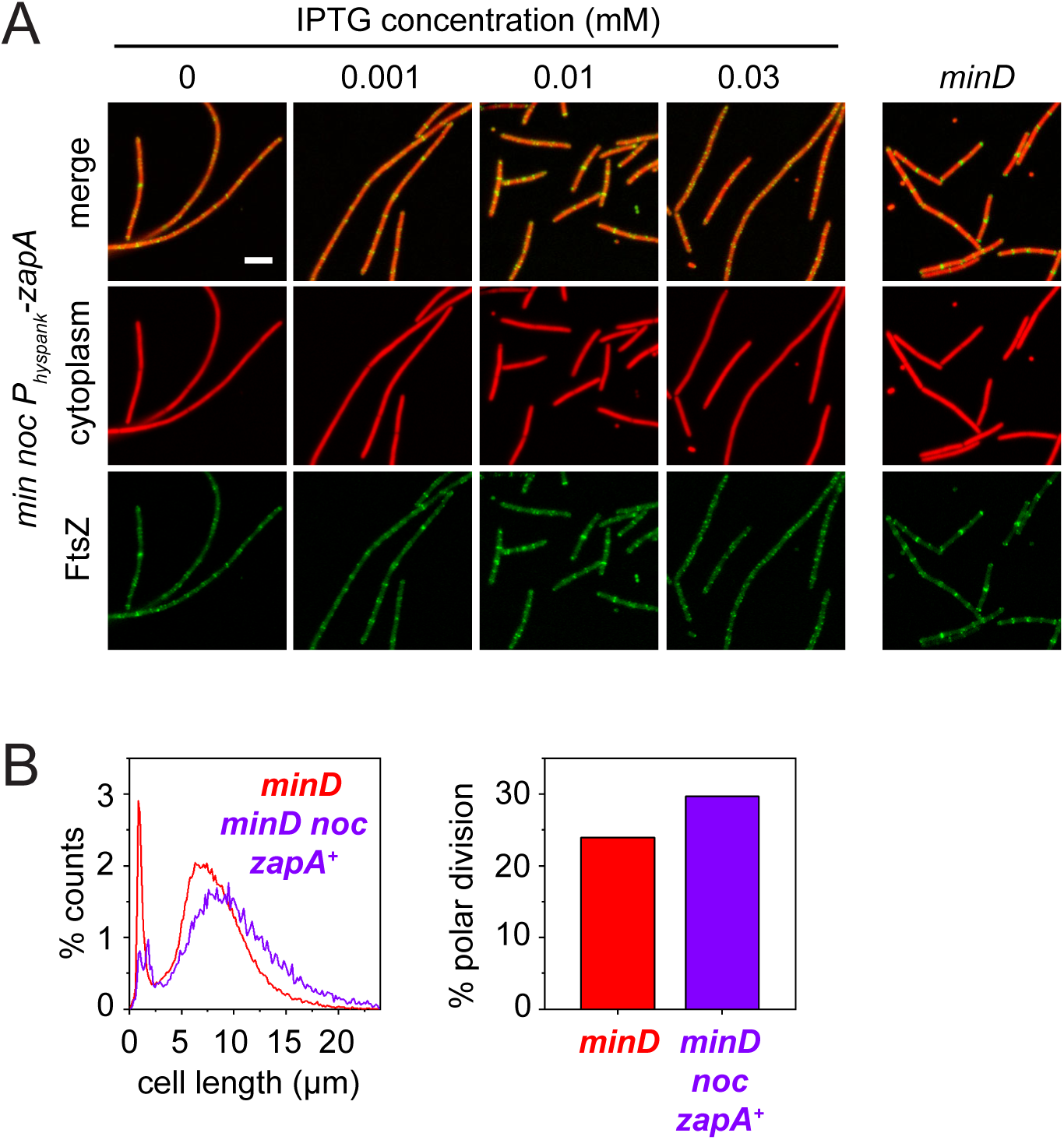
Artificial overexpression of ZapA suppresses the absence of Noc. A) Fluorescence images of a *minD noc* double mutant (DK8480) containing an IPTG-inducible copy of ZapA grown in broth culture containing the indicated amount of IPTG at 37°C. A *minD* mutant (DK8352) is provided for comparison. Constitutively expressed cytoplasmic mRFPmars false colored red, and mNeongreen-FtsZ false colored green. Scale bar is 5 μm and applies to all panels. B) Histogram of individual cell lengths for *minD* mutant (DK5155, blue) and a *minD noc* double mutant (DK8480, magenta) containing an IPTG-inducible copy of ZapA grown in 0.01 mM IPTG (*zapA^+^*) and measured on the microfluidic device. C) Bar graph of the percentage of cell division events that occurred on the cell pole for the *minD* mutant (DK5155, blue) and a *minD noc* double mutant (DK8480, magenta) containing an IPTG-inducible copy of ZapA grown in 0.01mM IPTG (*zapA^+^*). Manual counting of > 500 cell division events was performed to ensure minicells were accurately recorded.

ZapA expression might increase in the absence of ClpX and YjbH if the transcription factor Spx, normally degraded by the complex, instead accumulated and activated the expression of the *zapA* gene. To measure transcriptional regulation, the promoter region of *zapA* was cloned upstream of the *lacZ* gene encoding the LacZ β-galactosidase, and *P_zapA_-lacZ* expression increased two-fold when either *yjbH* or *clpX* was mutated (**Fig 3D**). We note that a two-fold increase in *zapA* transcription was consistent with the two-fold increase in ZapA protein level observed in Western blot (**Fig 3C**) and the two-fold increase in fluorescence intensity of ZapA-mNeongreen observed in either mutant background (**Fig 3A, right**). Moreover, the increase in transcription by reporter gene assay and increase in ZapA protein accumulation by Western blot analysis was abolished in both *clpX spx* and *yjbH spx* double mutants (**Fig 3C, 3D**). Finally, suppression by the *yjbH* mutation was Spx-dependent as a *minD noc yjbH spx* quadruple mutant was as temperature sensitive as the *minD noc* double mutant alone (**Fig 1D**). We note, however, that mutation of *spx* did not restore temperature sensitivity to the *minD noc clpX* triple mutant (**Fig 1D**). Nonetheless, we conclude that mutation of either *yjbH* or *clpX* overrides the absence of Noc by an Spx-dependent increase in ZapA expression. Furthermore, we infer that mutation of *clpX* confers an additional mechanism of growth rescue that is Spx-independent.

### An acceleration of cytokinesis overrides the absence of MinD in the *minD noc* double mutant

We wondered whether the Spx-dependent mechanism, as revealed by the *yjbH* mutation, also had an effect in the absence of MinD alone. One consequence of the *minD* mutation is that cell division events occur at the cell pole generating anucleoid minicells (**Fig 5A**),^52, 55, 87^ Mutation of *yjbH* reduced the polar division frequency of the *minD* mutant two-fold (**Fig 5A**), an effect likely mediated by elevated levels of Spx. Conversely, mutation of Spx exacerbated the *minD* minicell phenotype, by increasing the frequency of polar division two-fold (**Fig 5A**), such that nearly half of all divisions now occurred at the cell pole. Moreover, the mutation of *spx* was epistatic to *yjbH* as the polar division frequency of the *minD yjbH spx* triple mutant was elevated to levels that resembled the *minD spx* double mutant alone (**Fig 5A**). Thus, the frequency of minicell formation can be altered, and Spx appears to control an inhibitor of minicell formation. Finally, artificial overexpression of *zapA* in the *minD* mutant background also elevated minicell frequency (**Fig 5A**) suggesting that ZapA was likely not the inhibitor of minicell formation under Spx control. We infer that Spx regulates cell division not only by activating ZapA but also by controlling the expression or activity of another cell division regulator in parallel.

**Figure 5:**
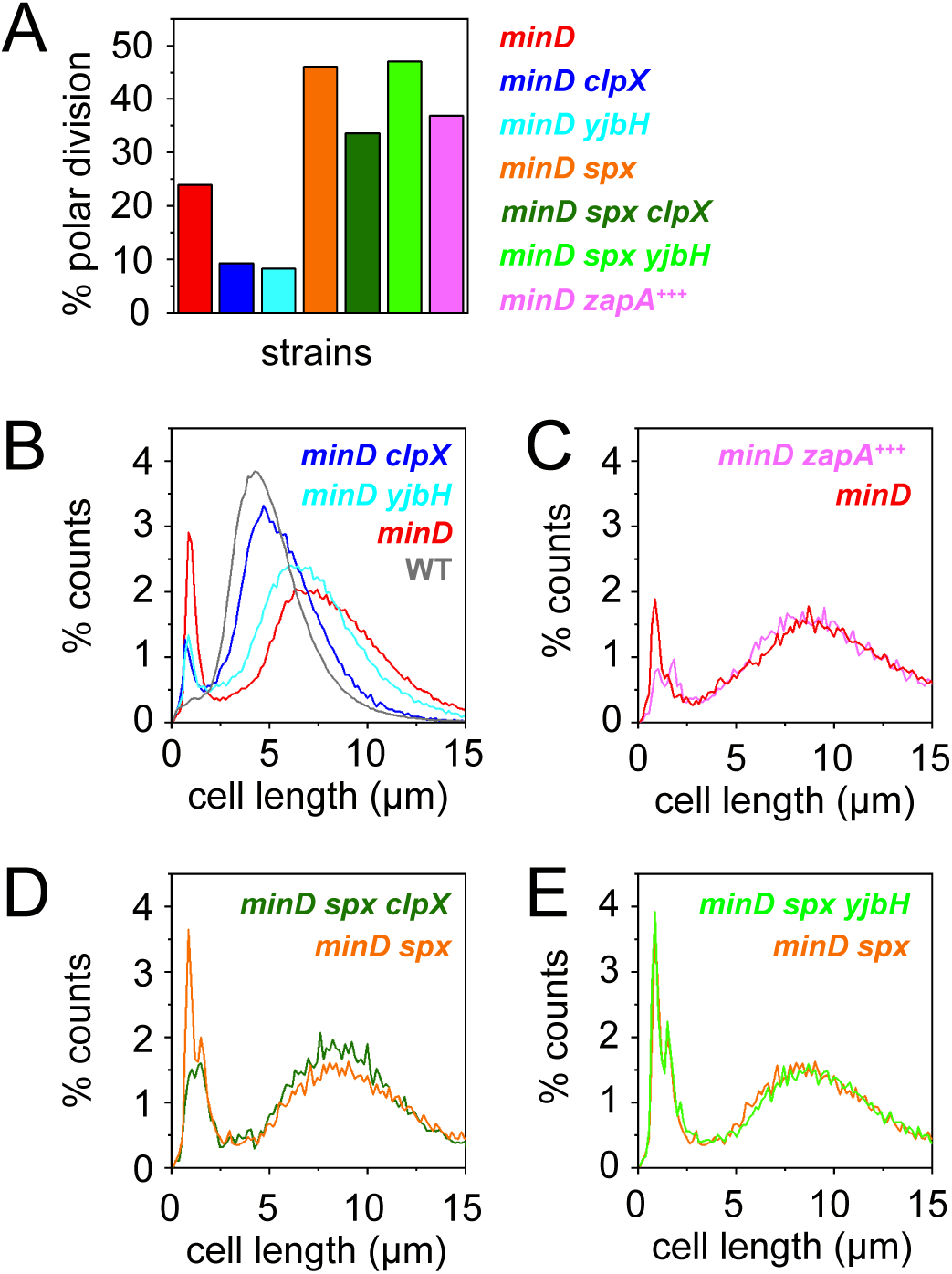
Minicell formation is inhibited by an Spx-dependent mechanism. A) Bar graph of the percentage of cell division events that occurred at the cell pole in the following mutants: *minD* (DK5155, red), *minD clpX* (DK7332, blue), *minD yjbH* (DK8747, cyan), *minD spx* (DK8918, orange), *minD spX clpX* (DK8922, olive), *minD spX yjbH* (DK8958, green), and a *minD* (*P_hyspank_- zapA*) grown with 0.01 mM IPTG (DK8353, magenta).For panels B to E, manual counting of > 400 cell division events for each strain was performed to ensure minicells were accurately recorded. B, C, D, E) Histogram of individual cell lengths for indicated mutants labeled with the same color scheme as described in panel A. Panel B also includes data for the wild type (gray, DK5133)

To determine whether mutation of *clpX* had the same effect as mutation of *yjbH* in the absence of *minD*, we examined division in the *minD clpX* double mutant. Consistent with ClpX-dependent proteolysis of Spx, mutation of *clpX* phenocopied mutation of *yjbH* in decreasing the *minD* polar division frequency (**Fig 5A**). Mutation of Spx, however, was only partially epistatic to mutation of ClpX as the polar division frequency of the *minD clpX spx* triple mutant was only partially elevated (**Fig 5A**). Mutation of *minD* also causes nucleoid-containing mother cells to become elongated (**Fig 5B**),^52, 53^ but mutation of *clpX* decreased *minD* mother cell length to that resembling wild type (**Fig 5B**), whereas simultaneous mutation of *minD* and *yjbH* (**Fig 5B**), or *minD* and *zapA* overexpression did not (**Fig 5C**). We conclude that the phenotypes of mother cell elongation and polar division frequency are separable, such that changes to one does not necessarily result in changes to the other. Nonetheless, mutation of *spx* was fully epistatic for cell length in both the *minD clpX spx* and *minD yjbH spx* triple mutants (**Fig 5D, Fig 5E**), suggesting that the effect of ClpX on cell length control can be overridden by the absence of Spx. Regardless, we sought to use the cell length differential between the *minD clpX* and *minD yjbH* double mutants to explore the Spx-independent mechanism of *minD noc* suppression.

Mutation of ClpX was shown to suppress the cell division defect that occurs when MinC and MinD are overexpressed, and the mechanism was attributed to ClpX acting as a direct inhibitor of FtsZ polymerization.^88, 89^ To determine the effect of ClpX on FtsZ dynamics, *minD yjbH* and *minD clpX* double mutants containing mNeongreen-FtsZ and cytoplasmic red fluorescence were grown and compared in microfluidic channels. 100 cells of each strain were chosen at random through the course of growth, and four events were sequentially measured: the appearance of an FtsZ-ring, accumulation of FtsZ to peak intensity, cell division, and FtsZ-ring disappearance (**Fig 6A,B**). Data from tracking the four events were then used to calculate various parameters of cell division.^18, 51, 65^ Each of the cell division parameters in the *minD yjbH* mutant were similar to that reported for *minD* (**Fig 6C**), and thus, any differences in the *minD clpX* double mutant would likely reflect the action of the Spx-independent mechanism of *minD noc* suppression. Mutation of *clpX* did not appear to alter FtsZ dynamics in the *minD* double mutant as polar FtsZ-rings exhibited indefinite persistence (**Fig 6C**), and the FtsZ-ring maturation time was similar to that of *minD yjbH* (**Fig 6C**). Instead, mutation of ClpX resulted in a statistically significant reduction in cytokinesis (defined as the time between peak FtsZ ring fluorescence and cell division) likely mediated at the level of divisome components downstream of FtsZ in the septation pathway. Moreover, faster cytokinesis produced shorter cells which may also have caused the *clpX* mutant to display a positive value in the medial Z-ring delay time indicating that the new FtsZ ring formed in the same generation as the division event (**Fig 6C**). We conclude that the Spx-independent mechanism of *minD* suppression does not change the rate of Z-ring maturation but instead accelerates the activation and/or function of the divisome.

**Figure 6.**
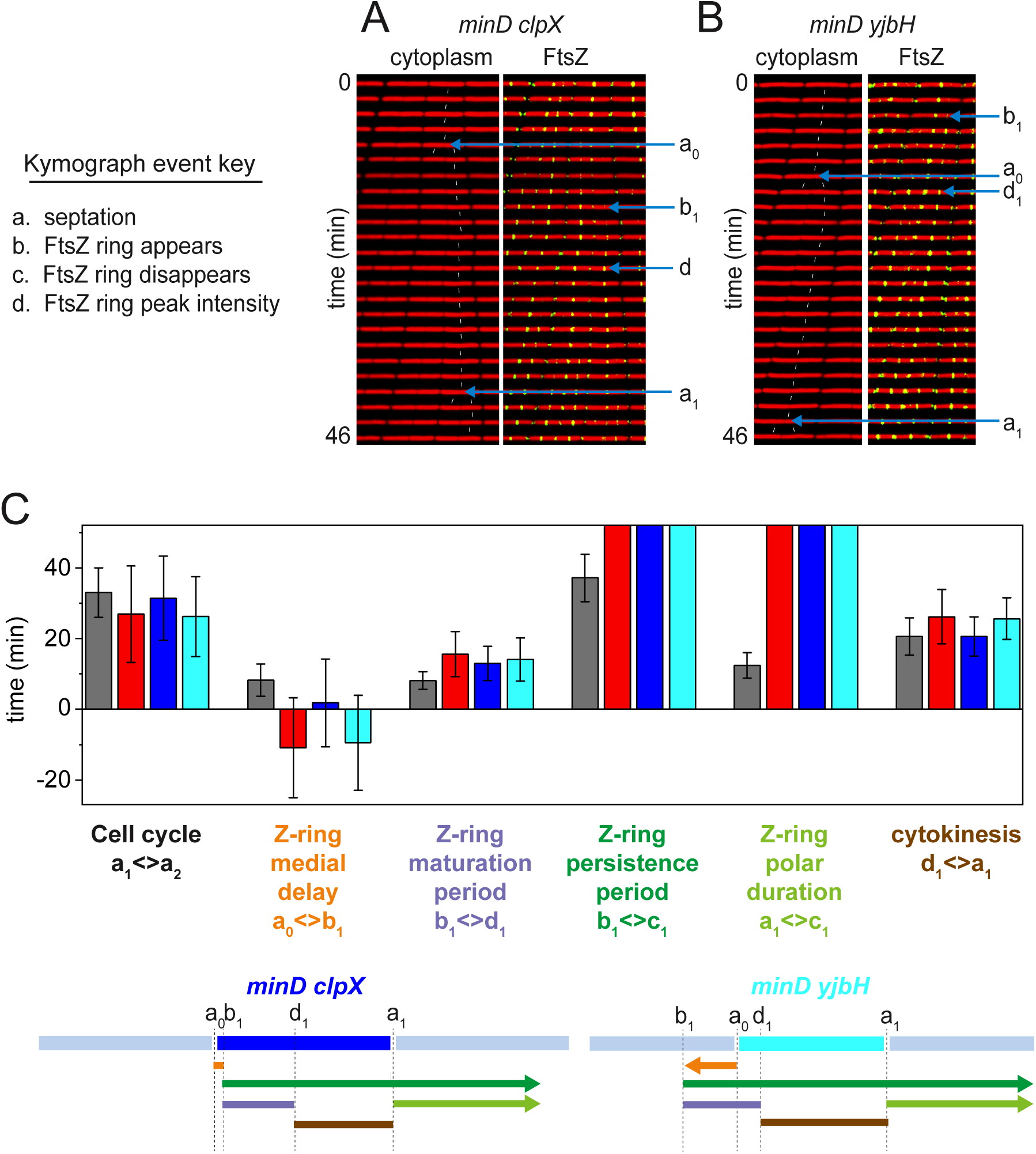
A Spx-independent mechanism reduces cytokinetic period for *minD* mutant. A and B) Kymograph analysis of the *minD clpX* (DK7332, A) and *minD yjbH* mutant (DK8747, B) grown in a microfluidic channel and imaged every 2 min. Cytoplasmic mCherry signal is false colored red (left) and overlaid with mNeongreen-FtsZ that is false colored green (right). Events necessary for defining division parameters are indicated and labeled a, septation; b, appearance of a nascent Z-ring; c, disappearance of a Z-ring; d, FtsZ peak intensity achieved. Each event designation is given a number: 0 for the preceding generation, 1 for the current generation, and 2 for the subsequent generation. Thin white lines indicate cell tracking and lineage analysis. C) Graphs of 100 manually tracked cell division cycles for the wild type (DK5133, gray), *minD* mutant (DK5155, red), *minD clpX* double mutant (DK7332, blue), and *min yjbH* double mutant (DK8747, cyan) presented as bars of average values and whiskers of standard deviation for the following parameters: “division time” is the time between septation events (between consecutive “a” events); “Z-ring medial delay” is the time between a septation event and the appearance of a Z-ring that will eventually give rise to the next medial division event (between an “a” event and a “b” event that will give rise to the next round of septation); “Z-ring maturation period” is the time between Z-ring appearance and when that Z-ring achieves peak local intensity (between consecutive “b” and “d” events); “Z-ring persistence period” is the time between the appearance of a Z-ring and the disappearance of that Z-ring (between consecutive “b” and “c” events); “Z-ring polar duration” is the time between a septation event and the disappearance of the Z-ring resulting from that septation event (between consecutive “a” and “c” events); and “cytokinesis period” is the time between a matured Z-ring to cell septation events (between a “d” event and the “a” event that is caused by that particular Z-ring). Note that the Z-ring medial delay of the *minD* and *minD yjbH* double mutants were negative on average because the medial Z-ring that would eventually promote cell division was formed in the preceding generation. The Z-ring persistence period and Z-ring polar duration in the *minD*, *minD clpX* and *minD yjbH* are indefinite because FtsZ persists at the division site after cell division and never disappears during the course of the experiments. D) Timelines of the various events, indicated in the bar graph, are color coded to match the indicated parameter of like color above, and annotated with relevant events marked by the defining letters.

## DISCUSSION

Min and Noc are thought to be two systems that help determine cell division site selection by controlling FtsZ localization. While neither system is essential in *B. subtilis*, cell division fails for unknown reasons when both are mutated simultaneously. Here, we show that in the absence of both Min and Noc, FtsZ becomes periodically enriched in inter-chromosomal spaces but fails to mature into an intense ring, such that cells subsequently elongate and eventually lyse. We interpret the double mutant phenotype in the context of models that suggest Noc sterically prevents FtsZ protofilaments from migrating away from the division plane, and Min depolymerizes FtsZ protofilaments to recycle monomers.^51, 65^ In the absence of Min, FtsZ monomers cannot be recycled, and FtsZ-ring maturation depends exclusively on the spatial concentration of de novo synthesized FtsZ by Noc corralling at the inter-chromosomal locations. In the absence of Noc, a subpopulation of protofilaments migrates from one division plane to the next, and the nascent Z-ring matures by Min-mediated monomer recycling. In the absence of both, the FtsZ protofilaments wander indefinitely and are neither stabilized by spatial restriction nor by sufficient monomer availability. Thus, neither system directs the localization of FtsZ to the inter-chromosome location per se, but both enable FtsZ to reach a sufficient local concentration needed to activate the divisome.

To better understand Z-ring maturation and activation of the divisome, we selected for second site suppressor mutations that restored high temperature growth in the absence of MinD and Noc. One class of suppressors disrupted components important for the regulatory proteolysis of the global stress response transcription factor Spx.^85, 86^ Spx is proteolyzed by the ClpXP machinery, and mutants that disrupted or impaired production of the Spx-specific adaptor protein YjbH restored growth.^82–84^ Consistent with the proteolytic model and suppression due to elevated levels of Spx, growth rescue by the absence of YjbH was abolished when Spx was mutated. Spx is a non-canonical DNA binding protein with a large regulon,^90–92^ but ZapA was predicted as a potentially important target because ZapA, but not FtsZ, protein levels were elevated in mutants enhanced for Spx. Moreover, Spx activated expression from the promoter of *zapA*, and previously published CHiP-seq analysis indicated that Spx binds upstream of the *P_zapA_* promoter.^90^ Thus, when YjbH is mutated, Spx levels increase in the cell, *zapA* transcription is increased, and growth is restored to the *minD noc* double mutant by increased levels of the ZapA protein.

Genetically, ZapA was both necessary and sufficient for growth rescue, as its mutation abolished suppression in the absence of YjbH, and its overexpression rescued growth to the *minD noc* double mutant background. ZapA is a small FtsZ-binding protein thought to condense and bundle protofilaments, and its artificial expression has been shown to suppress FtsZ condensation defects in the absence of Noc.^23, 65, 69, 70, 93^ Consistent with Noc suppression, titrated induction of ZapA gave rise to cell dimensions and FtsZ localization patterns in the *minD noc* double mutant that were indistinguishable from a *minD* mutant alone (**Fig 7**). The importance of ZapA however extends beyond the specific *yjbH* suppressor background as, unlike the absence of Spx, the absence of ZapA further lowered the permissive growth temperature for the *minD noc* double mutant from 30°C to 22°C. We note that transcription from the *P_zapA_* promoter increased to levels sufficient for suppression simply by growing cells at low temperature, but the elevated expression was only partly dependent on Spx (**Fig 3F**). Thus, we suggest that elevated *zapA* expression also helps explain the low temperature growth of *minD noc* double mutant. We infer that because Spx is not required for low temperature growth, other mechanisms of *zapA* activation are likely at work.

**Figure 7.**
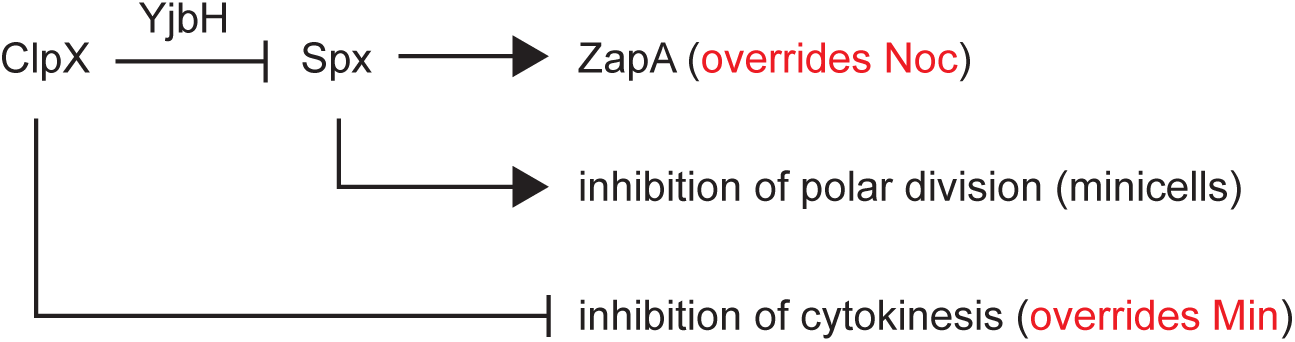
A model for the Spx-dependent and Spx-independent mechanisms of minD noc suppression. Arrows indicate activation, T bars indicate repression. Written in red are the interpretations of how the various pathways reported here override either the absence of Min or Noc.

We also found evidence of an Spx-independent mechanism that restored high-temperature growth to the *minD noc* double mutant, as mutation of ClpX did not precisely phenocopy mutation of YjbH. Specifically, whereas mutation of Spx abolished growth rescue in the absence of YjbH, Spx did not abolish rescue in the absence of ClpX (**Fig 1D**). Furthermore, mutation of ClpX also improved the temperature tolerance of the *minD noc zapA* triple mutant (**Fig 1C**). Thus, mutation of ClpX increases Spx and ZapA levels, but also engages an Spx- independent mechanism of suppression in parallel. Previous work indicated that ClpX interacts directly with FtsZ to inhibit FtsZ polymerization,^88, 89^ but our microfluidic analysis indicated that the Spx-independent suppression was not primarily mediated by changes in FtsZ dynamics. ClpX instead appeared to act downstream of FtsZ maturation. Indeed, the primary effect on division of the *minD clpX* double mutant was an acceleration of cytokinesis, likely by activating other components of the divisome (**Fig 7**). Our results may be consistent with the model of direct interaction between the two proteins, but whereas ClpX antagonizes FtsZ polymerization in vitro, the in vivo effect may primarily manifest at higher order FtsZ-divisome interactions. How ClpX antagonizes division is unknown, but we anticipate its further study may illuminate mechanisms of both FtsZ ring maturation and activation of the divisome.

Here, we report further evidence that the activity of the divisome is subject to regulation after FtsZ maturation. Specifically, the frequency of minicell formation could be increased or decreased two-fold when either Spx or YjbH was mutated, respectively. We infer that the frequency at which FtsZ rings become activated for cytokinesis is regulated, either across the entire cell, or specifically governed at the poles. A specific effect on polar divisome activation is implicated by the observation that the frequency of minicell formation is separable from cell length effects suggesting that medial and polar Z-rings are differentially regulated. If true, Z-rings may differ based on their location, an idea previously raised in *E. coli* based on differential sensitivity to inhibition by MinC/D.^94^ While the identity of the putative polar division inhibitor is unknown, our data suggest that inhibition is likely encoded as part of the Spx regulon, and we predict that this product may also be directly proteolyzed by ClpX (**Fig 7**). Why *B. subtilis* would encode a regulator of polar Z-ring activation during vegetative growth is unclear, but could be related to the process of sporulation in which polar septa give rise to the forespore under starvation conditions.^95, 96^

The results presented here indicate that cells mutated for both MinD and Noc fail to grow at high temperature due to a failure of Z-ring maturation, and that a variety of extragenic suppressor mutations restored growth by more than one mechanism. Two mechanisms were implicated by the study of ClpX and the stress response transcription factor Spx, but we note that mutations outside the system were also obtained. In particular, mutation of GgaB restored growth perhaps implicating an intersection between FtsZ maturation and the synthesis of teichoic acid. Furthermore, mutation of the transcriptional terminator Rho also restored growth, and strong phenotypes for *rho* mutations in *B. subtilis* have been notoriously difficult to identify.^97, 98^ Thus, in addition to the Spx-dependent and Spx-independent mechanisms of *minD noc* suppression reported here, there may be additional pathways revealed through the future study of the remaining suppressor classes. Whatever the case, the relative ease with which the conditional essentiality of the Min and Noc system can be bypassed suggest that there are likely multiple pathways that aid Z-ring maturation and may help explain why many bacteria lack either one or both systems. Thus, Min and Noc are simply two ways to help the Z-ring to mature rather than being integral parts of cell division site selection per se.

## MATERIALS AND METHODS

### Strains and growth conditions

*Bacillus subtilis* strains (**Table 2**) were grown in lysogeny broth (LB) (10 g tryptone, 5 g yeast extract, 10 g NaCl per liter) or on LB plates fortified with 1.5% Bacto agar at 37°C. Antibiotics were supplemented when appropriate at the following concentrations: 100 µg/ml spectinomycin, 5 µg/ml kanamycin, 10 µg/ml tetracycline, 5 µg/ml chloramphenicol, and macrolide-lincosamide-streptogramin B (MLS;1 µg/ml erythromycin, 25 µg/ml lincomycin). For *P_hyspank_* promoter-dependent gene expression, 1 mM Isopropyl-β-d-1-thiogalactopyranoside (IPTG) was added to the medium, if not otherwise indicated.

**Table 2:** Strains.

| Strain | Genotype |
| --- | --- |
| DK443 | *noc::kan* |
| DK1042 | Wild type |
| DK3391 | *amyE::P<sub>hysp</sub>ank-mRFPmars spec* |
| DK5133 | *mNeongreen-ftsZ ΔepsH amyE::P<sub>hysp</sub>ank-mcherry spec* |
| DK5155 | *mNeongreen-ftsZ ΔepsH amyE::P<sub>hysp</sub>ank-mcherry spec minD::TnYLB kan* |
| DK5565 | *clpX::kan* |
| DK5605 | *pTnFLX spec mls* |
| DK6330 | *amyE::P<sub>spank</sub>-noc spec* |
| DK6339 | *noc::mls* |
| DK6375 | *mNeongreen-ftsZ ΔepsH ycgO::P<sub>sigA</sub>-mRFP mars mls noc::kan amyE::P<sub>spank</sub>-noc spec* |
| DK6677 | *spx::spec* |
| DK6962 | *mNeongreen-ftsZ ΔepsH sacA::hbsu-mCherry cat noc::mls amyE::P<sub>spank</sub>-noc spec minD::TnYLB kan* |
| DK7279 | *minD::cat* |
| DK7298 | *noc::kan minD::cat* |
| DK7320 | *mNeongreen-ftsZ ΔepsH amyE::P<sub>hysp</sub>ank-mcherry spec clpX::kan* |
| DK7332 | *mNeongreen-ftsZ ΔepsH amyE::P<sub>hysp</sub>ank-mcherry spec clpX::kan minD::cat* |
| DK7369 | *noc::kan amyE::P<sub>spank</sub>-noc spec minD::cat* |
| DK7744 | *minD::cat* |
| DK8132 | *amyE::P<sub>hysp</sub>ank-zapA spec* |
| DK8138 | *ΔepsH zapA-mNeongreen amyE::P<sub>hysp</sub>ank-mcherry spec* |
| DK8140 | *noc::kan minD::cat amyE::P<sub>hysp</sub>ank-zapA spec* |
| DK8147 | *ΔepsH zapA-mNeongreen amyE::P<sub>hysp</sub>ank-mcherry spec clpX::kan* |
| DK8162 | *mNeongreen-ftsZ ΔepsH ycgO::P<sub>sigA</sub>-mRFP mars mls* |
| DK8174 | *ΔzapA* |
| DK8352 | *mNeongreen-ftsZ ΔepsH ycgO::P<sub>sigA</sub>-mRFPmars mls minD::cat* |
| DK8353 | *mNeongreen-ftsZ ΔepsH amyE::P<sub>hysp</sub>ank-zapA spec ycgO::P<sub>sigA</sub>-mRFPmars mls minD::cat* |
| DK8369 | *noc::mls clpX::kan minD::cat* |
| DK8387 | *ΔepsH zapA-mNeongreen amyE::P<sub>hysp</sub>ank-mCherry spec minD::cat clpX::kan noc::mls* |
| DK8390 | *mNeongreen-ftsZ ΔepsH amyE::P<sub>hysp</sub>ank-mCherry spec clpX::kan minD::cat noc::mls* |
| DK8394 | *ΔepsH zapA-mNeonGreen ycgO::P<sub>sigA</sub>-mRFPmars mls amyE::P<sub>spank</sub>-noc spec minD::cat* |
| DK8480 | *mNeongreen-ftsZ ΔepsH amyE::P<sub>hysp</sub>ank-zapA spec ycgO::P<sub>sigA</sub>-mRFPmars mls minD::cat noc::kan* |
| DK8524 | *mNeongreen-ftsZ amyE::P<sub>spank</sub>-noc spec ycgO::P<sub>sigA</sub>-mRFP mars mls* |
| DK8591 | *amyE::P<sub>zapA</sub>-lacZ cat* |
| DK8627 | *Spx::spec amyE::P<sub>zapA</sub>-lacZ cat* |
| DK8628 | *clpX::TnYLB kan amyE::P<sub>zapA</sub>-lacZ cat* |
| DK8631 | *Spx::spec clpX::TnYLB kan amyE::P<sub>zapA</sub>-lacZ cat* |
| DK8703 | *noc::kan minD::cat yjbH::TnFLX spec* |
| DK8704 | *noc::kan minD::cat yjbI::TnFLX spec* |
| DK8705 | *noc::kan minD::cat rho::TnFLX spec* |
| DK8706 | *noc::kan minD::cat upstream of yjbJ::TnFLX spec* |
| DK8707 | *noc::kan minD::cat rho::TnFLX spec* |
| DK8708 | *noc::kan minD::cat rho::TnFLX spec* |
| DK8709 | *noc::kan minD::cat upstream of clpP::TnFLX spec* |
| DK8710 | *noc::kan minD::cat ggaB::TnFLX spec* |
| DK8711 | *noc::kan minD::cat yaaT::TnFLX spec* |
| DK8712 | *noc::kan minD::cat clpX::TnFLX spec* |
| DK8713 | *noc::kan minD::cat hemX::TnFLX spec* |
| DK8714 | *noc::kan minD::cat ggaB::TnFLX spec* |
| DK8715 | *noc::kan minD::cat upstream of clpP::TnFLX spec* |
| DK8716 | *noc::kan minD::cat rho::TnFLX spec* |
| DK8738 | *noc::mls clpX::kan minD::cat spx::spec* |
| DK8741 | *mNeongreen-ftsZ ΔepsH amyE::P<sub>hysp</sub>ank-mcherry spec yjbH::TnKRM kan* |
| DK8742 | *ΔepsH zapA-mNeongreen amyE::P<sub>hysp</sub>ank-mcherry spec yjbH::TnKRM kan* |
| DK8747 | *mNeongreen-ftsZ ΔepsH amyE::P<sub>hysp</sub>ank-mcherry spec yjbH::TnKRM kan minD::cat* |
| DK8749 | *ΔepsH zapA-mNeongreen amyE::P<sub>hysp</sub>ank-mcherry spec noc::mls yjbH::TnKRM kan* |
| DK8767 | *spx::spec minD::cat noc::kan* |
| DK8800 | *ΔzapA minD::cat noc::kan* |
| DK8811 | *minD::cat yjbH::TnKRM kan noc::mls* |
| DK8817 | *mNeongreen-ftsZ ΔepsH amyE::P<sub>hyspank</sub>-mcherry spec noc::mls yjbH::TnKRM kan* |
| DK8824 | *ΔzapA noc::mls clpX::kan minD::cat* |
| DK8849 | *minD::cat yjbH::TnKRM kan noc::mls spx::spec* |
| DK8851 | *ΔzapA noc::mls yjbH::TnKRM kan minD::cat* |
| DK8880 | *clpX::kan spx::spec* |
| DK8881 | *yjbH::TnKRM kan spx::spec* |
| DK8883 | *yjbH::TnKRM kan amyE::P<sub>zapA</sub>-lacZ cat* |
| DK8890 | *spx::spec yjbH::TnKRM kan amyE::P<sub>zapA</sub>-lacZ cat* |
| DK8918 | *mNeongreen-ftsZ ΔepsH ycgO::P<sub>sigA</sub>-mRFPmars mls minD::cat spx::spec* |
| DK8922 | *mNeongreen-ftsZ ΔepsH ycgO::P<sub>sigA</sub>-mRFPmars mls minD::cat clpX::kan spx::spec* |
| DK8958 | *mNeongreen-ftsZ ΔepsH ycgO::P<sub>sigA</sub>-mRFPmars mls minD::cat spx::spec yjbH::TnKRM kan* |
| DS2231 | *clpX::TnYLB kan* |
| DS3187 | *ΔminJ minD::TnYLB kan amyE::P<sub>hag</sub>-hag(T209C) spec* |
| DS4522 | *spx::spec* |
| DS8875 | *yjbH::TnKRM kan* |

### Strain construction

All strains were generated by direct transformation of DK1042, a derivative of *B. subtilis* ancestral strain 3610 with enhanced frequency of natural competence for DNA uptake, or were transduced into DK1042 by SPP1-mediated transduction (**Table 2**).^99, 100^ All primers used for strain construction are listed in **Table S1**. Cells were mutated for the production of extracellular polysaccharide (EPS) to prevent biofilm formation within the microfluidic device by in-frame markerless deletion of *epsH* genes, encoding enzymes essential for EPS biosynthesis.^101, 102^ The mNeongreen fluorescent fusion to FtsZ or ZapA, a generous gift of Ethan Garner (Harvard University), was crossed into the indicated genetic background by SPP1 phage-mediated transduction, and the antibiotic resistance cassette was eliminated by cre-lox recombination.^21, 103^ The HBsu-mCherry fusion^67, 68^ was transduced into the indicated genetic backgrounds by SPP1 phage-mediated transduction. The *P_hyspank_-mcherry* inducible construct was introduced by integrating pEV6 at the *amyE* locus and selection for spectinomycin resistance.^104^ *<u>noc::kan</u>* <u>and *noc::mls*</u>. The *noc::kan* and the *noc::mls* insertion deletion alleles were generated with a modified “Gibson” isothermal assembly protocol.^105^ Briefly, the region upstream of *noc* was PCR amplified with the primer pair 3399/3400, and the region downstream of *noc* was PCR amplified with the primer pair 3401/3402. DNA containing a kanamycin resistance gene (pDG780)^106^ or erythromycin resistance gene (pAH52)^107^ were amplified with universal primers 3250/3251. The three DNA fragments were combined at equimolar amounts to a total volume of 5 µL and added to a 15 µl aliquot of prepared master mix (see below). The reaction mixture was incubated for 60 min at 50°C. The completed reaction was then PCR amplified with primers 3399/3402 to amplify the assembled product. The amplified product was transformed into competent cells of PY79 and then transferred to the 3610 background with SPP1-mediated generalized transduction. Insertions were verified by PCR amplification with primers 3399/3402.

#### amyE::P_hyspank_-mRFPmars

To generate the IPTG-inducible construct for mRFPmars expression, the gene encoding mRFPmars was PCR amplified from pmRFPmars (Addgene) with primers 4572/4573. The PCR product was purified, digested with NheI and SphI, and cloned into the NheI and SphI sites of pDR111 containing the *P_hyspank_* promoter, the gene conducing the LacI repressor protein, and an antibiotic resistance cassette between the arms of the *amyE* gene (generous gift from David Rudner, Harvard Medical School) to generate pDP430. The pDP430 plasmid was transformed into DK1042 to create DK3391.

#### ycgO::P_sigA_-mRFPmars

The constitutive mRFPmars construct was generated by isothermal assembly and direct transformation of the linear fragment into *B. subtilis*. The fragments upstream and downstream of *ycgO* were PCR amplified with primers 5270/5271 and 5272/5273, respectively, with chromosomal DNA from strain DK1042 as a template. The fragment containing the *P_sigA_* promoter was PCR amplified with primers 5268/5269 and chromosomal DNA from strain DK1042 as a template. The fragment containing mRFPmars and the spectinomycin resistance cassette was PCR amplified with primers 5266/5267 and chromosomal DNA from strain DK3391 as a template. The four PCR products were purified and mixed in an isothermal assembly reaction that was subsequently amplified by PCR with primers 5270/5273.

#### amyE::P_hyspank_-zapA spec

To generate the IPTG inducible construct for ZapA expression, the gene encoding ZapA was PCR amplified with primers 7278/7279 and chromosomal DNA from strain 3610 as a template. The PCR product was purified, digested with HindIII and SalI, and cloned into the HindIII and SalI sites of pDR111 to generate pYY8. The pYY8 plasmid was transformed into PY79 to create DK8132.

#### minD:: TnYLB kan

The *minD* gene was mutated by SPP1 phage-mediated transduction of a transposon mutant allele from DS3187 (*minD::TnYLB-1 kan* transposon insertion in the middle of the *minD* gene at sequence tag TAAATAGAG) reported previously.^40^

#### clpX::kan

The *clpX::kan* insertion deletion allele was generated with a modified “Gibson” isothermal assembly protocol. Briefly, the region upstream of *clpX* was PCR amplified with the primers 6048/6049, and the region downstream of *clpX* was PCR amplified with the primers 6050/6051. DNA containing the kanamycin resistance cassette (pDG780)^106^ was PCR amplified with universal primers 3250/3251. The three DNA fragments were combined following the “Gibson” isothermal assembly protocol above (see *noc::kan*). The assembled product was transformed into competent cells of PY79 to generate DK5565, and then transferred to the appropriate strain backgrounds by SPP1-mediated generalized transduction.

#### amyE::P_spank_-noc spec

To generate the IPTG inducible construct for Noc expression, the *noc* gene encoding was PCR amplified with primers 6409/6410 and chromosomal DNA from strain 3610 as a template. The PCR product was purified, digested with NheI and SalI, and cloned into the NheI and SalI sites of pDR110 containing the *P_spank_* promoter, the gene conducing the LacI repressor protein, and an antibiotic resistance cassette between the arms of the *amyE* gene (generous gift from David Rudner, Harvard Medical School) to generate pYY3. The pYY3 plasmid was transformed into PY79 to create DK6330.

#### minD::cat

The *minD::cat* insertion deletion allele was generated with a modified “Gibson” isothermal assembly protocol. Briefly, the region upstream of *minD* was PCR amplified with the primers 6739/6740, and the region downstream of *minD* was PCR amplified with the primers 6741/6742. DNA containing the chloramphenicol resistance cassette (pAC225)^107^ was PCR amplified with universal primers 3250/3251. The three DNA fragments were combined following the “Gibson” isothermal assembly protocol above (see *noc::kan*). The assembled product was transformed into competent cells of PY79 to generate DK7279, and then transferred to the appropriate strain backgrounds by SPP1-mediated generalized transduction.

#### ΔzapA

The in-frame deletion of *zapA* was generated by transducing the *zapA::kan* allele (BKK28610) obtained from the Bacillus Genetic Stock Center (The Ohio State University) into a strain of interest. The kanamycin cassette is flanked on either site by lox sequences and was evicted by transformation with the Cre-recombinase expressing plasmid pDR244 at 30°C and selection for spectinomycin resistance.^103^ Plasmid pDR244 was subsequently evicted by growth at 37°C in the absence of selection. Eviction of the kanamycin resistance cassette was confirmed by kanamycin sensitivity.

#### spx::spec

The *spx::spec* allele was the generous gift of Dr. John Helmann (Cornell University).

#### yjbH::TnKRM kan

The *yjbH::TnKRM kan* allele was previously reported.^108^

#### amyE::P_zapA_-lacZ cat

To generate the *P_zapA_-lacZ* transcriptional reporter plasmid pDP537, the promoter region of *zapA* (*P_zapA_*) was amplified from DK1042 chromosomal DNA with primers 7408/7049, digested with EcoRI and BamHI and cloned into the EcoRI/BamHI sites of pDG268 containing a chloramphenicol antibiotic resistance cassette and a polylinker upstream of the lacZ gene between two arms of amyE.^109^

### Isothermal assembly reaction

First a 5X ITA stock mixture was generated (500 mM Tris-HCL (pH 7.5), 50 mM MgCl_2_, 50 mM DTT (Bio-Rad), 31.25 mM PEG-8000 (Fisher Scientific), 5.02 mM NAD (Sigma Aldrich), and 1 mM of each dNTP (New England BioLabs)), aliquoted and stored at -80°C. An assembly master mixture was made by combining prepared 5X isothermal assembly reaction buffer (131 mM Tris-HCl, 13.1 mM MgCl_2_, 13.1 mM DTT, 8.21 mM PEG-8000, 1.32 mM NAD, and 0.26 mM each dNTP) with Phusion DNA polymerase (New England BioLabs) (0.033 units/µL), T5 exonuclease diluted 1:5 with 5X reaction buffer (New England BioLabs) (0.01 units/µL), Taq DNA ligase (New England BioLabs) (5328 units/µL), and additional dNTPs (267 µM). The master mix was aliquoted as 15 µl and stored at -80°C. DNA fragments were combined at equimolar amounts to a total volume of 5 µL and added to a 15 µl aliquot of prepared master mix. The reaction was incubated for 60 min at 50°C.

### SPP1 phage transduction

To 0.1 ml of dense culture grown in TY broth (LB broth supplemented after autoclaving with 10 mM MgSO_4_ and 100 µM MnSO_4_), serial dilutions of SPP1 phage stock were added and statically incubated for 15 min at 37°C. To each mixture, 3 ml TYSA (molten TY supplemented with 0.5% agar) was added, poured atop fresh TY plates, and incubated at 37°C overnight. Top agar from the plate containing near confluent plaques was harvested by scraping into a 50 ml conical tube, vortexed, and centrifuged at 5,000 x g for 10 min. The supernatant was treated with 25 µg/ml DNase final concentration before being passed through a 0.45 µm syringe filter and stored at 4°C. Recipient cells were grown to stationary phase in 2 ml TY broth at 37°C. 0.9 ml cells were mixed with 5 µl of SPP1 donor phage stock. 9 ml of TY broth was added to the mixture and allowed to stand at 37°C for 30 min. The transduction mixture was then centrifuged at 5,000 x g for 10 min, the supernatant was discarded, and the pellet was resuspended in the remaining volume. 100 µl of cell suspension was then plated on TY fortified with 1.5% agar, the appropriate antibiotic, and 10 mM sodium citrate.

### Microfluidic system

The microfluidic device was fabricated through a combination of electron beam lithography, contact photolithography, and polymer casting.^110^ Briefly, the microfluidic device is comprised of fluid and control layers both cast in poly(dimethylsiloxane) (PDMS) and a glass coverslip. The fluid layer lies between the control layer and glass coverslip and contains the microchannel array to trap the bacteria. Media and cells are pumped through the microfluidic channels by on-chip valves and peristaltic pumps that are controlled pneumatically through the top control layer. Each pneumatic valve is controlled by software to apply either vacuum (0.3 bar) or pressure (1.3 bar) to open or close individual valves, respectively.

### On-device cell culture

Before cells were loaded into the microfluidic device, the microchannels were coated with 1% bovine serum albumin (BSA) in LB medium for 1 h to act as a passivation layer. Then, all the channels were filled with LB medium containing 0.1% BSA. A saturated culture of cells (∼10 μL) was added through the cell reservoir and pumped into the cell-trapping region. During cell loading, vacuum was applied to the control layer to lift up the microchannel array. After a sufficient number of cells were pumped underneath the channel array, positive pressure was applied to trap individual cells in those channels. Medium was pumped through the microchannels to flush away excess cells and maintain steady-state cell growth in the channel array.

### Time-lapse image acquisition

After inoculation in the microfluidic channels, a period of roughly 3 h elapsed during which cells adjusted to the growth conditions, and steady-state cell growth was maintained and monitored over the next 10 h. Fluorescence microscopy was performed either on a Nikon Eclipse Ti-E microscope or an Olympus IX83 microscope. The Nikon Eclipse Ti-E microscope was equipped with a 100× Plan Apo lambda, phase-contrast, 1.45 numerical aperture (NA) oil immersion objective and a Photometrics Prime95B sCMOS camera with Nikon Elements software (Nikon, Inc.). Fluorescence signals from mCherry, mRFPmars and mNeongreen were captured from a Lumencor SpectraX light engine with matched mCherry, mCherry, and yellow fluorescent protein (YFP) filter sets, respectively, from Chroma. The Olympus IX83 microscope was equipped with an Olympus UApo N 100×/1.49 oil objective and a Hamamatsu EM-CCD digital camera operated with MetaMorph Advanced software. Fluorescence signals from mCherry, mRFPmars, and mNeongreen were excited with an Olympus U-HGLGPS fluorescence light source with matched tetramethyl rhodamine isocyanate (TRITC), TRITC, and fluorescein isothiocyanate (FITC) filters, respectively, from Semrock. Images were captured from at least eight fields of view at 2-min intervals, if not otherwise indicated. The channel array was maintained at 37°C with a TC-1-100s temperature controller (Bioscience Tools). For all direct comparisons, the same microscope and settings were used.

### Data analysis

An adaptation period following exposure to illumination was observed; thus, data analysis was restricted to periods of steady state. Cell identification and tracking were analyzed by MATLAB programs (The MathWorks, Inc.).^110^ The program extracted fluorescence intensity along a line profile down the longitudinal center of each microchannel in the array. The cytoplasmic mCherry or mRFPmars line profiles showed a flat-topped peak on the line where a cell was located, and a local 20% decrease in fluorescence intensity was used to identify cell boundaries after division. Division events were conservatively measured as the time at which one cell became two according to the decrease in fluorescence intensity. Moreover, cell bodies were tracked from frame to frame in order to construct lineages of cell division, and cell body intensity was determined by the integration of the cytoplasmic mCherry or mRFPmars signal intensities within the confines of the cell. Signals from HBsu-mCherry, mNeongreen-FtsZ and ZapA-mNeongreen were similarly tracked and measured along the length of the cell. FtsZ and ZapA line profiles were normalized by cell body intensity in order to minimize intensity differences among frames and across different fields of view.

### Microscopic image acquisition on agarose pads

Cells were grown to mid-log phase in liquid culture at 37°C and then imaged on 1% agarose pads in 1×PBS. Fluorescence microscopy was performed with a Nikon 80i microscope with a Nikon Plan Apo 100× phase-contrast objective and an Excite 120 metal halide lamp. Cytoplasmic mRFPmars was visualized with a C-FL HYQ Texas Red filter cube, and mNeongreen-FtsZ and ZapA-mNeongreen were visualized with a C-FL HYQ FITC filter cube. Images were captured with a Photometrics Coolsnap HQ2 camera in black and white, false colored, and superimposed with Metamorph image software.

### Western blotting

*B. subtilis* strains were grown in LB medium to an optical density at 600 nm (OD_600_) of ∼1.0, and 1 mL was harvested by centrifugation, resuspended to 10 OD_600_ in lysis buffer (20 mM Tris [pH 7.0], 10 mM EDTA, 1 mg/mL lysozyme, 10 μg/mL DNAse I, 100 μg/mL RNAse I, 1 mM phenylmethylsulfonyl fluoride [PMSF]), and incubated 30 min at 37°C. Ten microliters of lysate was mixed with 2 μl 6× SDS loading dye. Samples were separated by 15% sodium dodecyl sulfate-polyacrylamide gel electrophoresis (SDS-PAGE). The proteins were electroblotted onto nitrocellulose and developed with a 1:80,000 dilution of anti-SigA primary antibody (generous gift of Masaya Fujita, University of Houston), a 80,000 dilution of anti-FtsZ primary antibody (generous gift of Petra Levin, Washington University), a 1:1000 dilution of anti- ZapA primary antibody (generous gift from David Rudner, Harvard Medical School), and a 1:1000 dilution secondary antibody (horseradish peroxidase [HRP]-conjugated goat anti-rabbit immunoglobulin G). Immunoblot was developed with the Immun-Star HRP developer kit (Bio-Rad).

### Transposon mutagenesis

The transposon plasmid was introduced into the *min noc* double mutant (DK7298) of *B. subtilis* by SPP1 phage transduction at room temperature, selecting for MLS resistance, and incubated at room temperature for 72 h. Subsequently, cells were incubated for 16 h at room temperature in LB medium. To obtain individual colonies, the cell culture was diluted 1:10 with LB medium, and 100 µl of the dilution was plated on a LB plate fortified with 1.5% agar, supplemented with spectinomycin, and grown at the non-permissive temperature (42°C) overnight. To confirm that the transposon was linked to the suppressor mutation, a lysate was generated on the suppressor mutant, and the transposon was transduced to the parent *min noc* double mutant strain (DK7298) lacking the suppressor selecting for spectinomycin resistance and grown at 42°C.

### Transposon insertion site identification by IPCR

Cells were grown in LB medium at 37°C for 6 h. DNA was isolated, and 6 µg was digested with 2U/µg DNA of the restriction endonuclease Sau3AI (NEB). Ligation with 400 U of T4 DNA ligase (NEB) was performed at room temperature for 2 h with 1.5 µg of the digested DNA. Inverse PCR (IPCR) was performed with 0.5 µg of ligated DNA with oligonucleotides 6212 and 6420 in a standard PCR program with a 3 min extension time on a Mastercycler thermocycler (Eppendorf). Phusion DNA polymerase (NEB) was used for PCR amplification. The amplified fragment was analyzed by Sanger sequencing (IDT) with the universal sequencing oligonucleotide 6224.

### β-Galactosidase assay

One milliliter of cells was harvested from a mid-log-phase (OD_600_ ∼0.5) culture grown in LB broth shaken at 37°C and resuspended in an equal volume of Z buffer (40 mM NaH_2_PO_4_, 60 mM Na_2_HPO_4_, 1 mM MgSO_4_, 10 mM KCl and 38 mM β-mercaptoethanol). To each sample, lysozyme was added to a final concentration of 0.2 mg/ml and incubated for 15 min at 30°C. Each sample was diluted appropriately to 500 µl in Z buffer, and the reaction was started with 100 µl of 4 mg/ml O-nitrophenyl β-D-galactopyranoside (in Z buffer) and stopped with 250 µl of 1M Na_2_CO_3_. The OD_420_ of the reaction mixtures was recorded, and the β-galactosidase-specific activity was calculated according to the equation: [OD_420_/(time×OD_600_)] ×dilution factor ×1000.

### Dilution plating

*B. subtilis* cells were grown in LB medium at room temperature to mid-log phase and then diluted to an optical density at 600 nm (OD_600_) of 1.0. The diluted cultures were serially diluted (10^-1^ to 10^-6^), 10 µl of each dilution was spotted onto an LB plate, and were grown overnight at either 30°C or 37°C.

## ACKNOWLEDGEMENTS

We thank Jinsheng Zhou, David Kysela, Sampriti Mukherjee, and Stephen Olney for technical support. We thank Ethan Garner for the ZapA-mNeongreen and mNeongreen-FtsZ fusions, Xindan Wang for the *sacA::hbsu-mCherry cat* fusion, David Rudner for the anti-ZapA primary antibody, Petra Levin for the anti-FtsZ primary antibody, Masaya Fujita for the anti-SigA primary antibody, and the IU Nanoscale Characterization Facility for use of its instruments. The work was supported in part by FAPESP grant 16/05203-5 to FJGH, NIH grant R01 GM113172 to SCJ, and NIH grant R35 GM131783 to DBK.

## FIGURE LEGENDS

**Figure S1:**
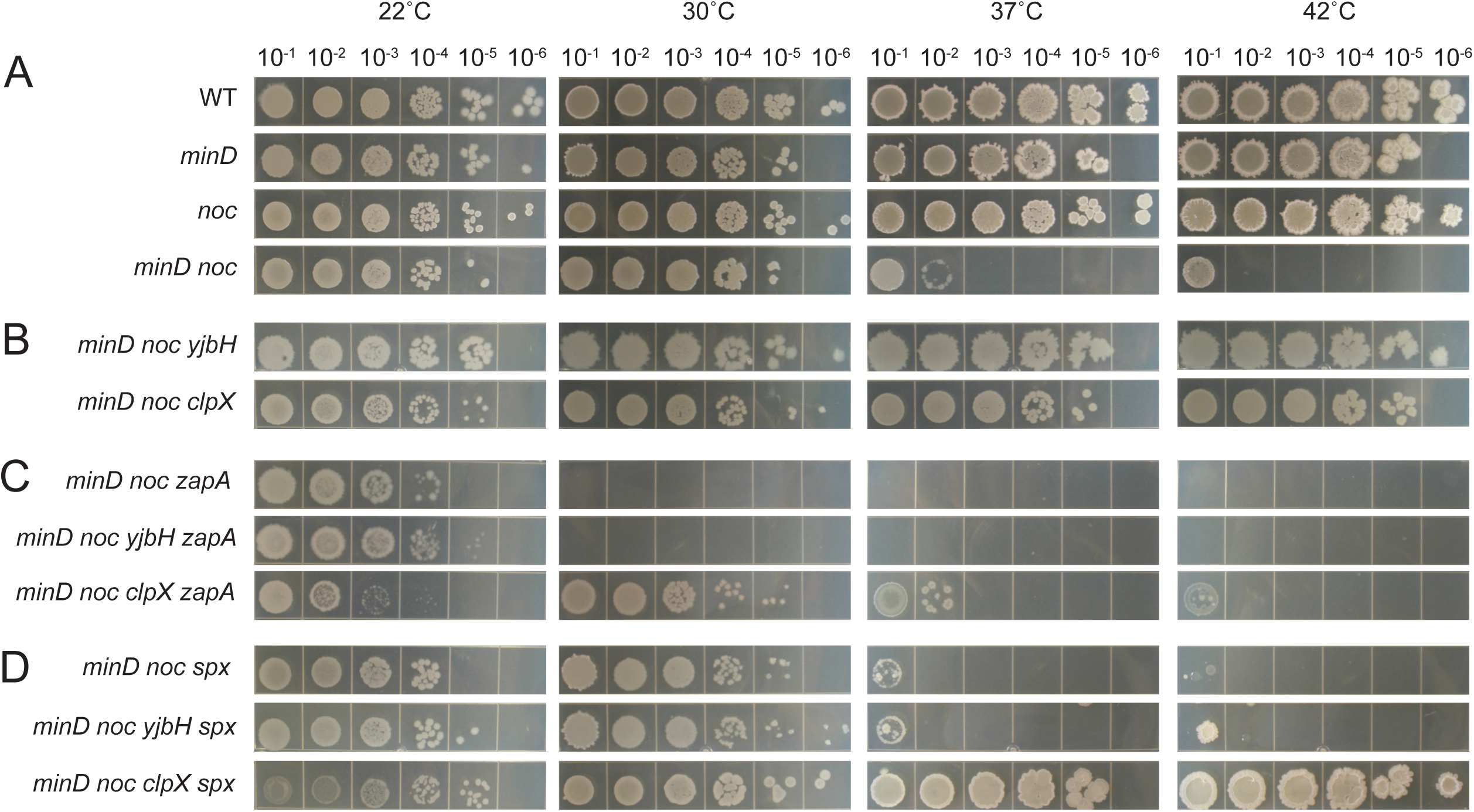
y*j*bH restores high-temperature viability to the *minD noc* double mutant through *spx*. Dilution plating of A) the wild type (DK1042), *minD* (DK7744), *noc* (DK443), *minD noc* (DK7298), B) *minD noc clpX* (DK8369), *minD noc yjbH* (DK8811), C) *minD noc zapA* (DK8800), *minD noc zapA yjbH* (DK8851), *minD noc zapA clpX* (DK8824), and D) *minD noc spx* (DK8767), *minD noc spx yjbH* (DK8849), *minD noc spx clpX* (DK8738) at 22°C, 30°C, 37°C, and 42°C. Exponential growing liquid culture were diluted to OD_600_ ≈ 1, and then serially diluted to 10^-1^, 10^- 2^, 10^-3^, 10^-4^, 10^-5^, and 10^-6^. 10 µL of cells were spotted for each concentration.

**Figure S2:**
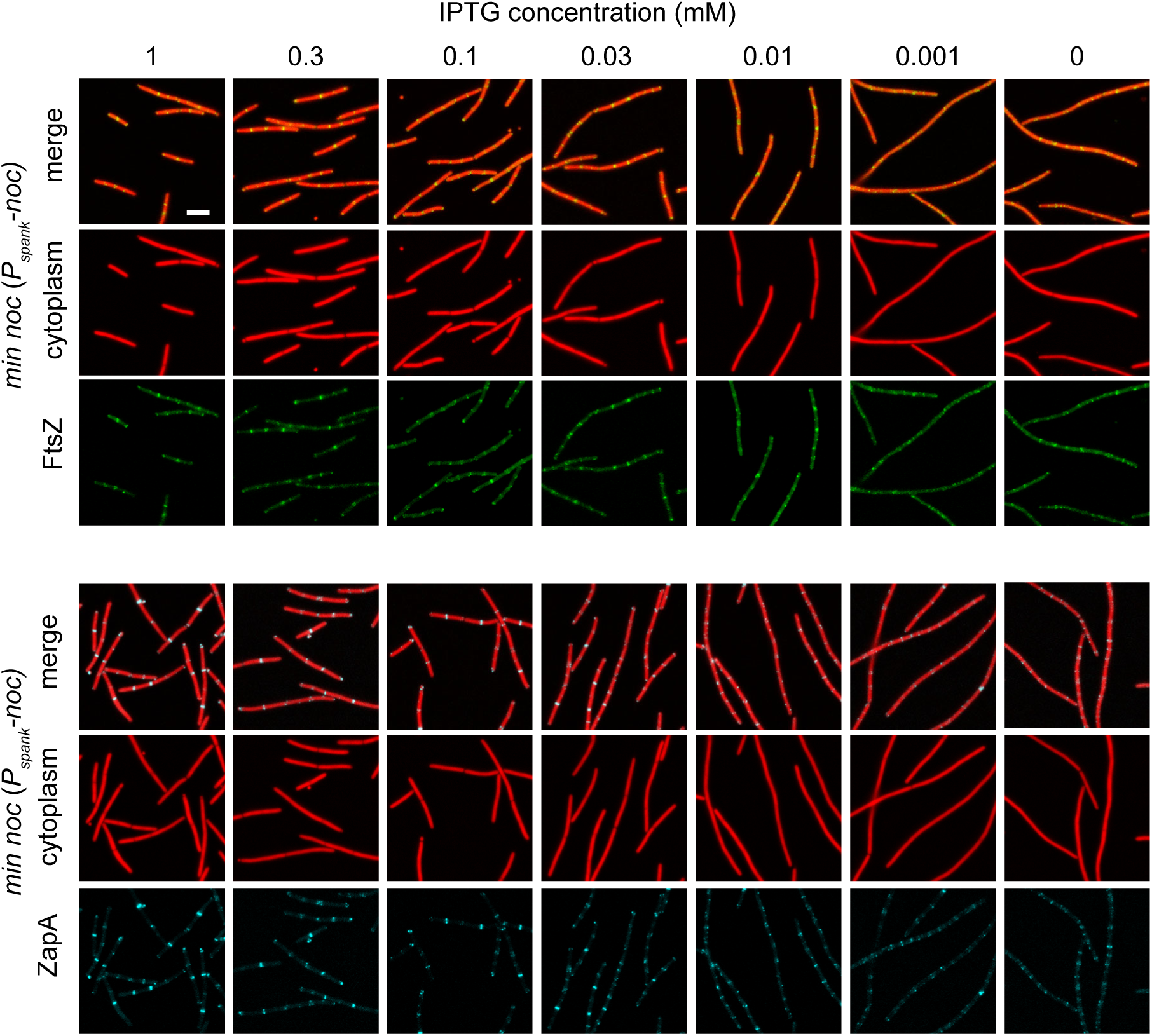
Overexpression of Noc from *P_spank_-noc* in the *minD no*c double mutants. Fluorescence images of the *min noc* double mutants (top, DK6375; bottom, DK8394) containing an IPTG-inducible copy of Noc (*P_spank_-noc*) grown in broth culture with the indicated amount of IPTG. Constitutively expressed cytoplasmic mRFPmars false colored red, mNeongreen-FtsZ false colored green, and ZapA-mNeongreen false colored cyan. Scale bar is 5 μm and applies to all panels.

